# The “replacing surgery” of cpDNA: *de novo* chemical synthesis and *in vivo* functional testing of *Chlamydomonas* chloroplast genome

**DOI:** 10.1101/2023.01.05.522807

**Authors:** Chunli Guo, Guiying Zhang, Hui Wang, Rui Mei, Xinyi Li, Hui Li, Bin Jia, Chaogang Wang, Zhangli Hu

**Affiliations:** Guangdong Technology Research Center for Marine Algal Bioengineering, Guangdong Provincial Key Laboratory for Plant Epigenetics, College of Life Sciences and Oceanography, Shenzhen University, Shenzhen 518055, China; College of Physics and Optoelectronic Engineering, Shenzhen University, Shenzhen 518060, China; Shenzhen Engineering Laboratory for Marine Algal Biotechnology, Longhua Innovation Institute for Biotechnology, Shenzhen University, Shenzhen 518055, China

**Keywords:** Chloroplast genome (cpDNA), SynCpV1.0, Functional testing, Replacing cpDNA, Photosynthetic efficiency, *Chlamydomonas reinhardtii*

## Abstract

We have successfully designed and synthesized the 221,372-bp cpDNA SynCpV1.0 with the native cpDNA of *Chlamydomonas reinhardtii* as the template. Homoplasmic SynCpv1.0-harboring algal strains were obtained by biolistic transformation and selected with an ascending gradient of antibiotic pressure. Meanwhile, we were pleasantly surprised to find that SynCpV1.0 was able to re-introduce and replicate normally after the total DNA of transplastomic algal strains were transformed to *Escherichia coli*, it indicated that SynCpV1.0 was able to shuttle between *C. reinhardtii* and *E. coli*. Finally, we analyzed the photosynthetic properties of SynCpV1.0-harboring transplastomic strains, the results showed that they exhibited the same photosynthetic efficiency as the wild strain of *C. reinhardtii* CC125, and could rescue the photosynthetic defect in mutant strain of *C. reinhardtii* CC5168. Herein, we have performed the “replacing surgery” of cpDNA and established an ideal platform to complete multiple cycles of “Design-Build-Test” for optimizing the cpDNA of photosynthetic organisms.

**Highlight:**

- An artificial cpDNA SynCpV1.0 is constructed by *de novo* chemical synthesis.
- The “replacing surgery” of cpDNA was performed in the chloroplast of *C. reinhardtii*
- It is found that artificial cpDNA was able to shuttle between *Chlamydomonas* chloroplast and *E. coli*.
- Establish an ideal platform to complete multiple cycles of “Design-Build-Test” for optimizing the cpDNA.

**One-Sentence Summary:** The chloroplast genome can be replaced by a complete synthesized genome and performs the designed biological function in *C. reinhardtii*.

## INTRODUCTION

Photoautotrophic organisms such as plants, algae always provide oxygen and food that is necessary for the ecosystem on earth, and play an important role on reducing carbon dioxide emissions, preventing climate change and addressing the global agricultural crisis(Bailey-Serres et al., 2019; Wheeler and von Braun, 2013). Meanwhile, chloroplasts are the most important photosynthetic organelle in higher plants and algae, in which light energy is converted into chemical energy by synthesizing organic matters from carbon dioxide. Although genome of chloroplast (cpDNA) can encode most of the key genes, which are involved in photosynthesis, it is difficult for conventional breeding methods to simultaneously overcome the many bottlenecks that restrict photosynthetic efficiency due to the low probability of spontaneous mutations in these genes of native cpDNA (Ort et al., 2015). Genetic engineering and synthetic biology are required to achieve various goals, including the manipulation of photosynthetic components for more efficient light harvesting and energy conversion(Perera-Castro and Flexas, 2020), “upgradation” of carbon fixation pathways(Simkin et al., 2017; Simkin et al., 2015) elimination of ubiquitous key factors affecting photosynthetic efficiency, e.g., the low catalytic efficiency of Rubisco enzyme(Cummins et al., 2018; Sharwood, 2017) and synthesis of high-value pharmaceutical and food additive products using chloroplasts (Dyo and Purton, 2018). Therefore, *de novo* total synthesis, *in vitro* assembly, and functionally characterization of synthetic cpDNA are prerequisites for the rational design and reconstruction of photosynthetic systems in organisms.

Synthetic genomics refers to the *de novo* chemical synthesis of the entire genome or most parts of the genome *via* a series of technical approaches. Synthetic genomics began with the chemical synthesis of yeast alanine tRNA gene(Agarwal et al., 1970), which was followed by the artificial modifications and synthesis of various viral genomes, e.g., poliovirus (Cello et al., 2002), φX174 bacteriophage(Smith et al., 2003), T7 bacteriophage(Chan et al., 2005), severe acute respiratory syndrome-like coronavirus(Becker et al., 2008), and West Nile Virus(Orlinger et al., 2010). The most representative works of synthetic genomics including the synthesis and minimization of *Mycoplasma mycoides* genome, the recoding of the *Escherichia coli* genome, and the artificial synthesis of *Saccharomyces cerevisiae* chromosomes have been previously described(Hutchison et al., 2016; Ostrov et al., 2016). However, the development of *de novo* chemical synthesis of the chloroplast genome is relatively slow. Higher plants are obligate photoautotrophs, which restrict the artificial modification in their photosynthetic systems. Still we have not found a big story related to artificial modification in photosynthetic systems in higher plant except study on the complete cpDNA of *Oryza sativa* which was assembled from the chemically synthesized DNA fragments in *Bacillus subtilis*(Itaya et al., 2008). However, the authors did not perform functional testing on the synthetic cpDNA in *O. sativa*. In 2012, the assembly of *C. reinhardtii* cpDNA in yeast was reported using several large DNA fragments from BAC libraries of *C. reinhardtii* cpDNA, it allowed simultaneous modifications of multiple sites within the cpDNA of *C. reinhardtii*(O’Neill et al., 2012). However, it has not yet been achieved to date for the *de novo* chemical synthesis and *in vivo* functionally characterization of synthetic cpDNA in photosynthetic organisms.

In this study, we aimed to design and construct an artificial cpDNA, and transformed it into chloroplast of *C. reinhardtii* for replacing the original cpDNA and exerting its biological functions.

## RESULTS

### Strategy for design, synthesis and assembly of SynCpV1.0

The SynCpV1.0 was designed by using the native cpDNA of *C. reinhardtii* (Accession no: PMC5775909, 205.535kb) as the template and inserting BAC backbone, *aphVIII, aadA*, and multiple HA-tags. The cpDNA of *C. reinhardtii* has a GC content of about 34% divided by two ~22-kb inverted repeats (IRa and IRb) into two single-copy regions of ~80 kb. In addition, the cpDNA contains short dispersed repeats (SDRs) that account for more than 20% of the sequence (Gallaher et al., 2018; Maul et al., 2002). In this study, the strategy for synthesis, assembly, and functional assessment of the synthetic cpDNA (SynCpV1.0) was carried out according to the principle of the DBT cycle in *C. reinhardtii* (Figure 1).

**Figure 1.**
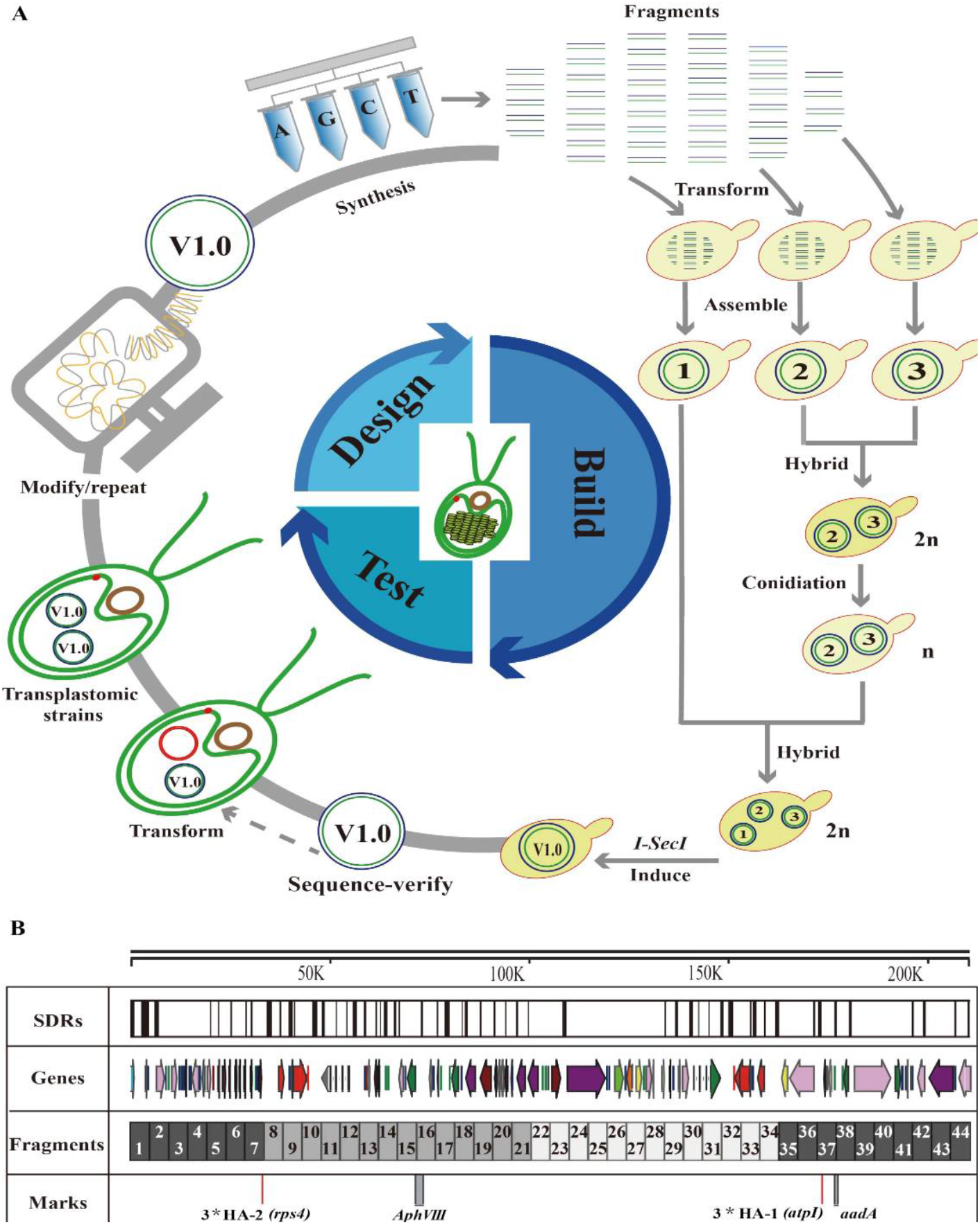
Strategy design, synthesis, assembly and functional test of SynCpV1.0. (A) The schematic diagram of the steps in the global synthesis of chloroplast genome in *C. reinhardtii* from synthetic oligonucleotides. Including redesign of the genome sequence *via* computer, *de novo* chemical synthesis, assemble in *S. cerevisiae*, transfer into *E. coli*, transfer into chloroplast of *C. reinhardtii*, and testing the function of SynCpV1.0. (B) Design of the chloroplast genome of *C. reinhardtii*. The length of cpDNA SynCpV1.0 is 221,372-bp, and was divided into 44 primary fragments, in which *aphVIII*, *aadA*, and multiple HA-tags were inserted. About 20% of the chloroplast genome of *C. reinhardtii* is repetitive DNA containing various types of SDRs in the intergenic regions.

Here, the antibiotic resistance markers, including *aphVIII* and *aadA*, and multiple HA-tags were inserted into the native cpDNA of *C. reinhardtii* to obtain SynCpV1.0. The designed SynCpV1.0, which has a total of 221,372 bp in length, was divided into 44 primary fragments (seg1–seg44, ~4.9 kb each) with 120-bp homologous flanking sequences at both ends. The *5’atpA--aphVIII--3’rbcL* expression cassette was located at 71,780–74,066, while the *5’atpA--aadA--3’rbcL* expression cassette was located at 176,191–177,820 in the SynCpV1.0. Additionally, the HA-tags were inserted into *atpI* on seg37 and *rsp4* on seg7, respectively. Subsequently, SynCpV1.0 was assembled in yeast. In brief, those 44 primary fragments were first assembled into three secondary fragments, i.e., chunk1, chunk2, and chunk3. Each of the secondary fragments was then inserted with a recognition site of *I-SceI*. After being transformed into the yeast, the expression of the endonuclease *I-SceI* was induced to linearize the secondary fragments, which were then self-assembled *via* homologous recombination into SynCpV1.0 (Figure 1).

### Assembly of SynCpV1.0 in *Saccharomyces cerevisiae*

The primary fragments ranging seg1–7 and seg35–seg44 were co-transformed and assembled into the secondary fragment chunk1 in *S. cerevisiae* BY4741, which was then screened using the synthetic complete medium lacking uracil (SC-URA medium). After that, the transformants carrying the accurately assembled chunk1 were identified *via* junction PCR (Supplementary figure S1A). The primary fragments ranging from seg7–seg21 were co-transformed into *S. cerevisiae* HWY175, which was then screened with the complete synthetic medium lacking L-leucine (SC-LEU medium). The transformants harboring the accurately assembled secondary fragment chunk2 were subsequently identified *via* junction PCR (Supplementary figure S1B). The primary fragments ranging from seg22–seg35 were co-transformed into *S. cerevisiae* BY4741. After screening using the SC-Met medium, the transformants carrying the accurately assembled secondary fragment chunk3 were identified *via* junction PCR (Supplementary figure S1C). Subsequently, the yeast transformants carrying chunk2 and chunk3 were hybridized and screened to obtain the auxotrophic haploid yeast strain harboring chunk2 and chunk3.

Following the hybridization with the yeast strain carrying chunk1, the hybrid yeast strain containing all three secondary fragments, i.e., chunk1, chunk2, and chunk3, were screened and transformed with the ZLP012 plasmid, which was constructed by adding *I-SecI* to pRS413 (-His) (Supplementary figure S2). After that, the expression of *I-SceI* was induced to linearize chunk1, chunk2, and chunk3 plasmids, which eventually self-assemble into SynCpV1.0 in the transformant. The yeast strain containing the complete SynCpV1.0 genome was then identified *via* junction PCR (Figure 2A). The total genomic DNA extracted from the yeast strain was subject to third-generation sequencing. The sequencing results showed that the sequencing reads covered the entire sequence of SynCpV1.0 genome designed in this study (Figure 2B), suggesting that we successfully generated the cpDNA of *C. reinhardtii via* complete chemical synthesis and assembly in the yeast assembly system.

**Figure 2.**
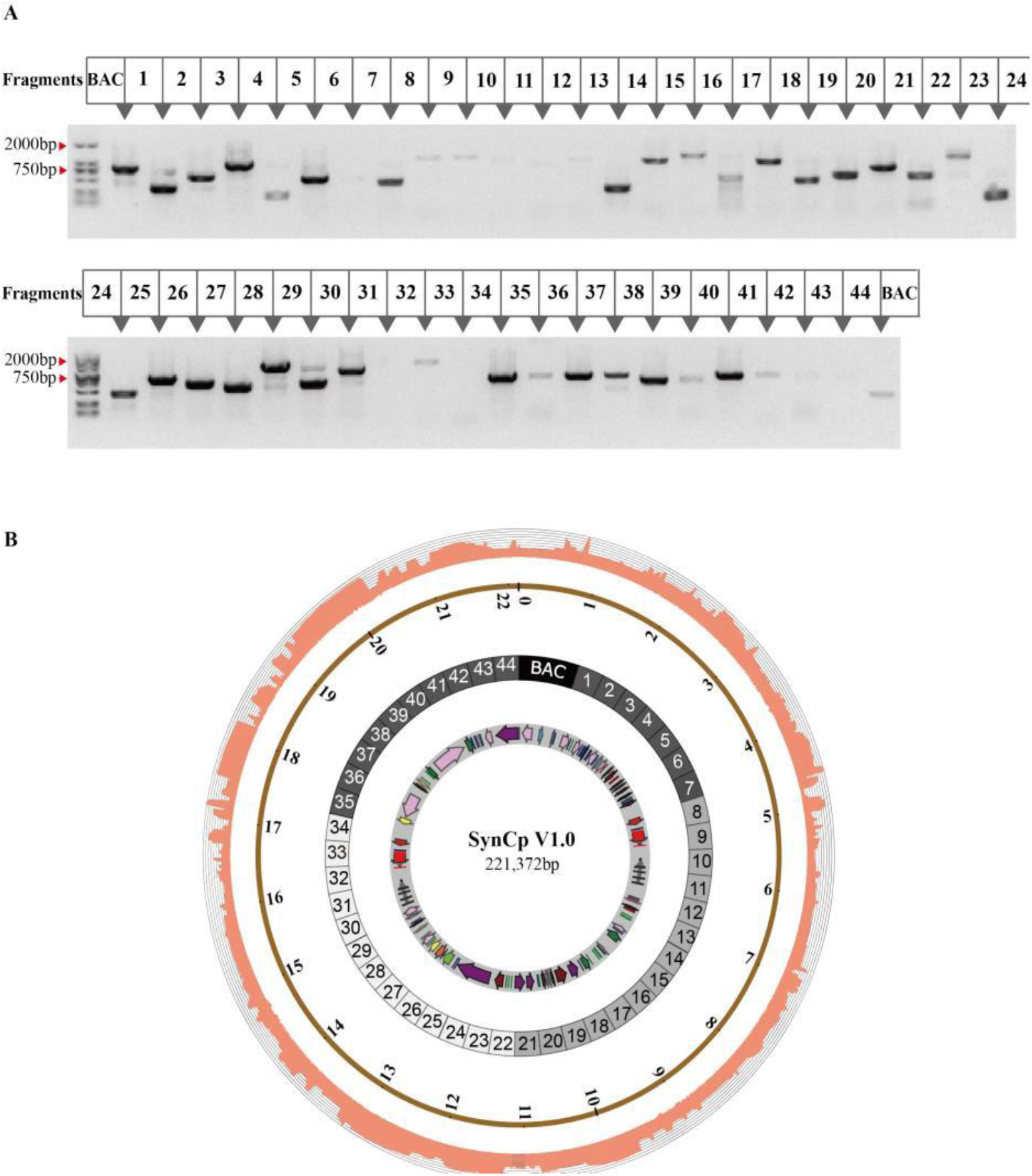
High-fidelity oligonucleotide assembly in yeast. (A) Analysis of the assembled synthetic plasmid carrying SynCpV1.0 *via* 44 PCR amplification for the adjacent fragments in yeast. PCR products were separated by gel-electrophoresis on a 1% agarose gel, DL2,000 marker was applied, and each triangle indicated the amplification product between two adjacent fragments. (B) Coverage of the synthetic plasmid sequence and the designed cpDNA sequence of *C. reinhardtii*. The synthetic plasmid sequence is consistent with designed cpDNA sequence.

### Transformation and homogenization of transplastomic algal strains

SynCpV1.0 were extracted from yeast, and then transformed into *E. coli* for self-replication and isolation of high-quality SynCpV1.0 plasmid. After that, SynCpV1.0 was transformed into the chloroplast of *C. reinhardtii* by the biolistic method. In this study, *C. reinhardtii* (CC125 and CC5168 (*ΔpsbH*)) were selected as the recipient strains for SynCpV1.0 transformation, after which the CC125 transformant was screened with the tris-acetate-phosphate (TAP) agar plate supplemented with 150 mg/L of streptomycin, while the CC5168 transformant was screened by the Sueoka’s high salt photoautotrophic medium. Statistical data showed that the genetic transformation assays yielded a total of 2,315 transplastomic strains, comprising 1,013 transplastomic strains derived from the CC125 recipient strain and 1,302 transplastomic strains from the CC5168 recipient strain. As exogenous sequences (e.g., plasmid vector backbone for assembly, resistance genes *aphVIII* and *aadA* for screening, as well as HA-tags) have been incorporated into SynCpV1.0, the transplastomic strains can be identified *via* PCR assays with primers targeting those exogenous sequences. Totally, we selected 14 transplastomic strains containing all the above exogenous gene fragments to carry out homogenization screening, of which 6 transplastomic strains were derived from *C. reinhardtii* CC125 and 8 transplastomic strains from *C. reinhardtii* CC5168 (Figures 3A).

**Figure 3.**
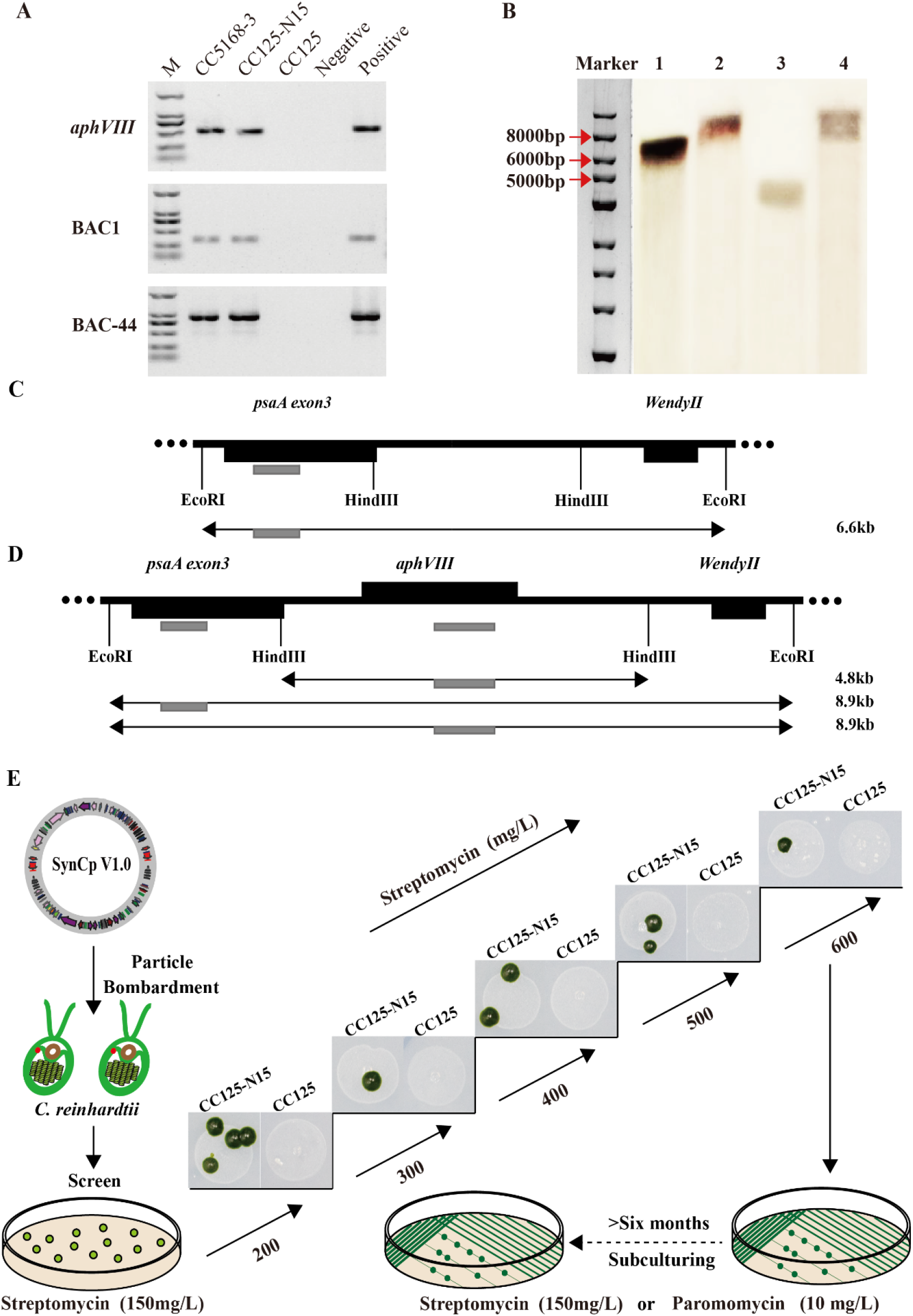
Screening and homogeneity verification for positive transplastomic strains carrying the SynCpV1.0. (A)Validation of the exogenous fragments. CC5168-3 and CC125-N15 are transplastomic strains carrying SynCp V1.0, wild type strain CC125 and H2O were used as negative controls. Positive control is the plasmid carrying SynCp V1.0, M represents DL2,000 marker. (B) RFLP-southern blotting analysis of positive transplastomic strains carrying the SynCpV1.0. Lane 1 represents the genomic DNA of *C. reinhardtii* CC125 digested with *EcoR* I and hybridized with the *psaA* probe. Lane 2 is transplastomic strain CC125-N15 digested with *Eco*R I and hybridized with the *psaA* probe. Lane 3 is genomic DNA of transplastomic strain CC125-N15 digested with *Hind* III, and subjected to Southern Blot with *aphVIII* probe. Lane 4 is genomic DNA of transplastomic strain CC125-N15 digested with *Eco*R I, and subjected to Southern Blot with *aphVIII* probe. (C) The genomic DNA of *C. reinhardtii* CC125 digested with *Eco*R I and hybridized with the *psaA* probe to obtain 6.6-kb hybrid fragment. Grey rectangle indicates the *psaA* probe.(D) The genomic DNA of *C. reinhardtii* CC125-N15 digested with *Eco*R I or *Hin*d III and hybridized with the *psaA* probe or *aphVIII* probe, to obtain 8.9-kb and 4.8-kb hybrid fragment respectively. Grey rectangle indicates the *psaA* probe or *aphVIII* probe. (E) Homogenization of transplastomic strains carrying the SynCpV1.0. The monoclonal transplastomic strains from selective medium with 150 mg/L streptomycin were transferred to fresh TAP medium with 200 mg/L streptomycin. Then monoclonal colonies were transferred to new medium with 300 mg/L streptomycin and the streptomycin concentration was gradually improve to 600 mg/L, finally to obtain the high degree of homogeneity of transplastomic strains which kept in selective medium with streptomycin or spectinomycin for more than 6 months. Wild type CC125 as control could not grow normally on every streptomycin medium.

To obtain a transplastomic *C. reinhardtii* strain completely homoplasmic for SynCpv1.0, those transplastomic strains were exposed to the selective pressure exerted by an ascending gradient of streptomycin ranging from 150 mg/L to 600 mg/L, and then subcultured with antibiotic medium (streptomycin or spectinomycin) for more than 6 months (Figures 3E).

To determine the homogeneity of SynCpV1.0 in the chloroplast of *C. reinhardtii*, the transplastomic strains were subjected to restriction fragment length polymorphism (RFLP) Southern blot analysis by probes targeting its endogenous gene *psaA* and exogenous gene *aphVIII*, respectively. The genomic DNAs of *C. reinhardtii* CC125 and its transplastomic strain CC125-N15 were digested with *Eco*RI and hybridized with the *psaA* probe. The results showed that *C. reinhardtii* CC125 and transplastomic strain CC125-N15 respectively, gave rise to hybridization bands of 6,681 bp and 8,968 bp, indicated that the presence of the *aphVIII* expression cassette in SynCpV1.0 (Figure 3B, C and D). Subsequently, the genomic DNA of transplastomic strain CC125-N15 was digested with *Eco*RI, and subjected to Southern Blot with *aphVIII* probe, which showed that hybridization bands was 8,968 as same as the bands from *psaA* probe (Figure 3B, C and D). Moreover, the genomic DNA of transplastomic strain CC125-N15 was digested with *HindIII*, and subjected to Southern Blot with *aphVIII* probe, which formed a band of 4,829 bp indicating that the *aphVIII* expression cassette is located between 79,599 bp and 88,566 bp within SynCpV1.0 (Figure 3B, C, and D) as the predesignated position in SynCpV1.0. The RFLP-Southern blot results showed that all transplastomic strains formed single hybridization bands, suggesting a high degree of homoplasmy whereby the fully synthetic SynCpV1.0 replaced the wild-type cpDNA in those transplastomic strains.

### Characterization of the *in vivo* pattern of SynCpV1.0

To characterize *in vivo* pattern of SynCpV1.0 in the chloroplast of *C.reinhardtii*, we carried out experiments on re-transforming SynCpV1.0 from transplastomic algal strains to *E.coli*. SynCpV1.0 comprises a backbone sequence that allows self-replication in *E. coli*. It should be capable of replication in *E. coli* if it remains circular after replacing the wild-type cpDNA in positive transplastomic algal strains. Therefore, the total genomic DNAs of *C. reinhardtii* CC125 and its transplastomic strain CC125-N15 were extracted and re-introduced respectively into *E. coli*. The transformation of the genomic DNA from the transplastomic strain CC125-N15 into *E. coli* yielded single colonies, which carry all 44 primary fragment (Figure 4), while the transformation of the genomic DNA from *C. reinhardtii* CC125 into *E. coli* did not produce any transformant.

**Figure 4.**
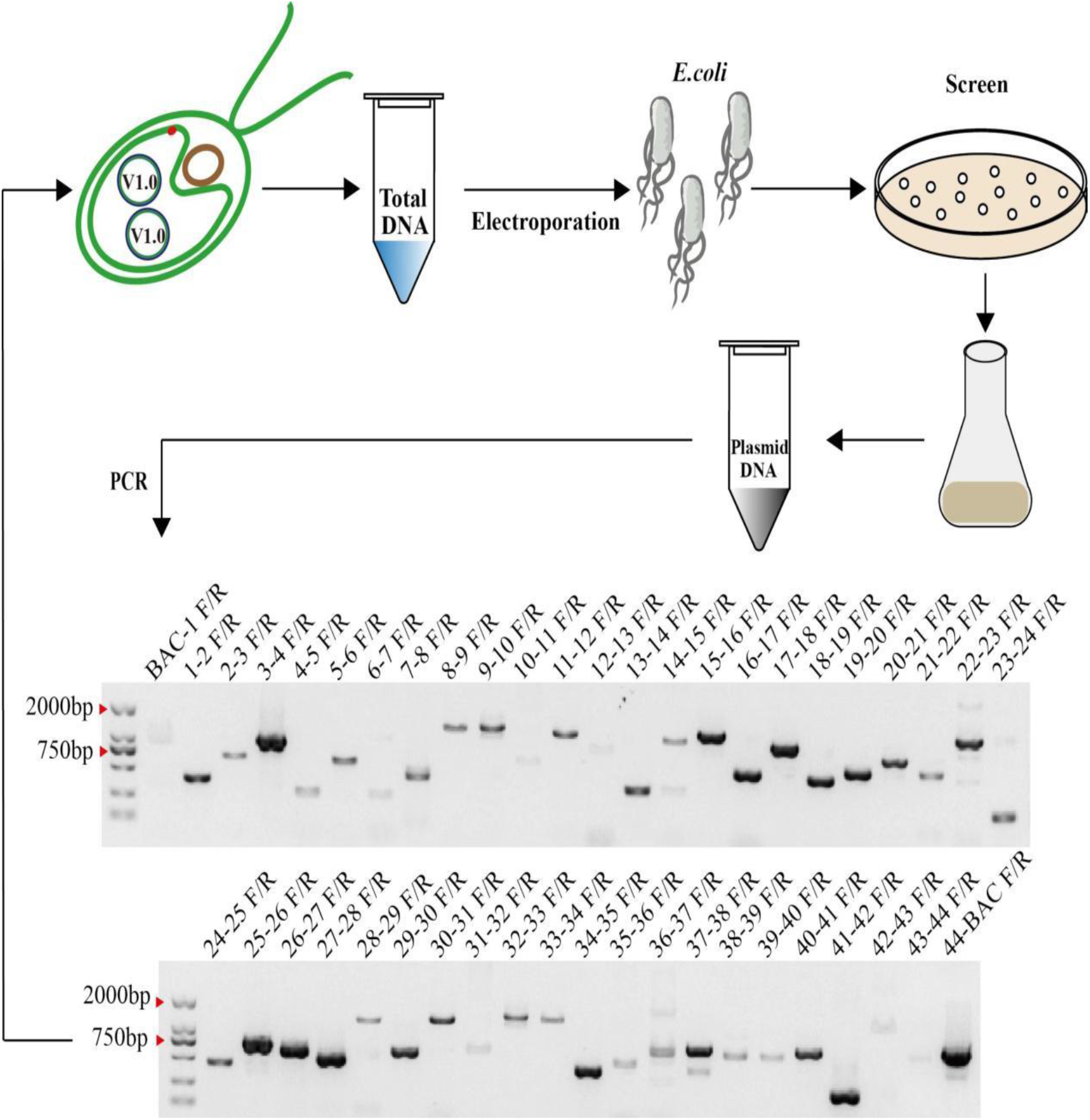
Re-transformation of SynCpV1.0 from transplastomic algal strains into *E. coli*. The total genomic DNA from transplastomic strains harboring SynCpV1.0 was extracted and transformed into *E. coli* EPI300 by electroporation method. The plasmids of positive clones were amplified with 45 junction PCR. BAC-1 and BAC-44 were the primers that targeted to the adjacent sequence between vector BAC and seg1, BAC and seg44, respectively. The lanes from 1-2F/R to 43-44F/R were the primers that amplified the fragments among the adjacent primary fragments. M represents DL 2,000 marker.

The above results implied that SynCpV1.0 might exist as a circular genomic DNA in transplastomic *C. reinhardtii* chloroplast, and the regulatory elements on its vector backbone enable SynCpV1.0 to replication in *E. coli*. Therefore, the fully chemically synthesized SynCpV1.0 can replicate in both *C. reinhardtii* chloroplast and *E. coli*. It is the first time to find that artificial cpDNA can shuttle between plant cells and *E. coli*. it will greatly facilitate the re-engineering and functional improvement of *C. reinhardtii* cpDNA. At present, gene editing of *Chlamydomonas* chloroplast is unsuccessful, our experiments provide a new breakthrough to optimize cpDNA using an efficient gene editing system of bacteria. SynCpV1.0 can be developed into a shuttle plasmid between the chloroplast of *C. reinhardtii* and *E. coli*. Herein, we have established an ideal platform to complete multiple cycles of “Design-Build-Test” for optimizing the cpDNA of photosynthetic organisms.

### Functional analysis of SynCpV1.0 in C. *reinhardtii*

The observations obtained from this study revealed that the growth performance of transplastomic strains was similar to that of the recipient algal strain, indicating that SynCpV1.0 can maintain the normal growth of algal cells. *C. reinhardtii* CC5168 is a *psbH*-deficient mutant strain incapable of photosynthesis activities and can only grow slowly in complete darkness compared with wide type. It reached only a very low cell density after 11 days of culture, while its transplastomic strains displayed a similar growth performance as observed in *C. reinhardtii* CC125, indicating that SynCpV1.0 incorporated in the *C. reinhardtii* CC5168, which replaced the mutant cpDNA with deletion of *psbH* and restored its photoautotrophy (Figure 5C and D).

**Figure 5.**
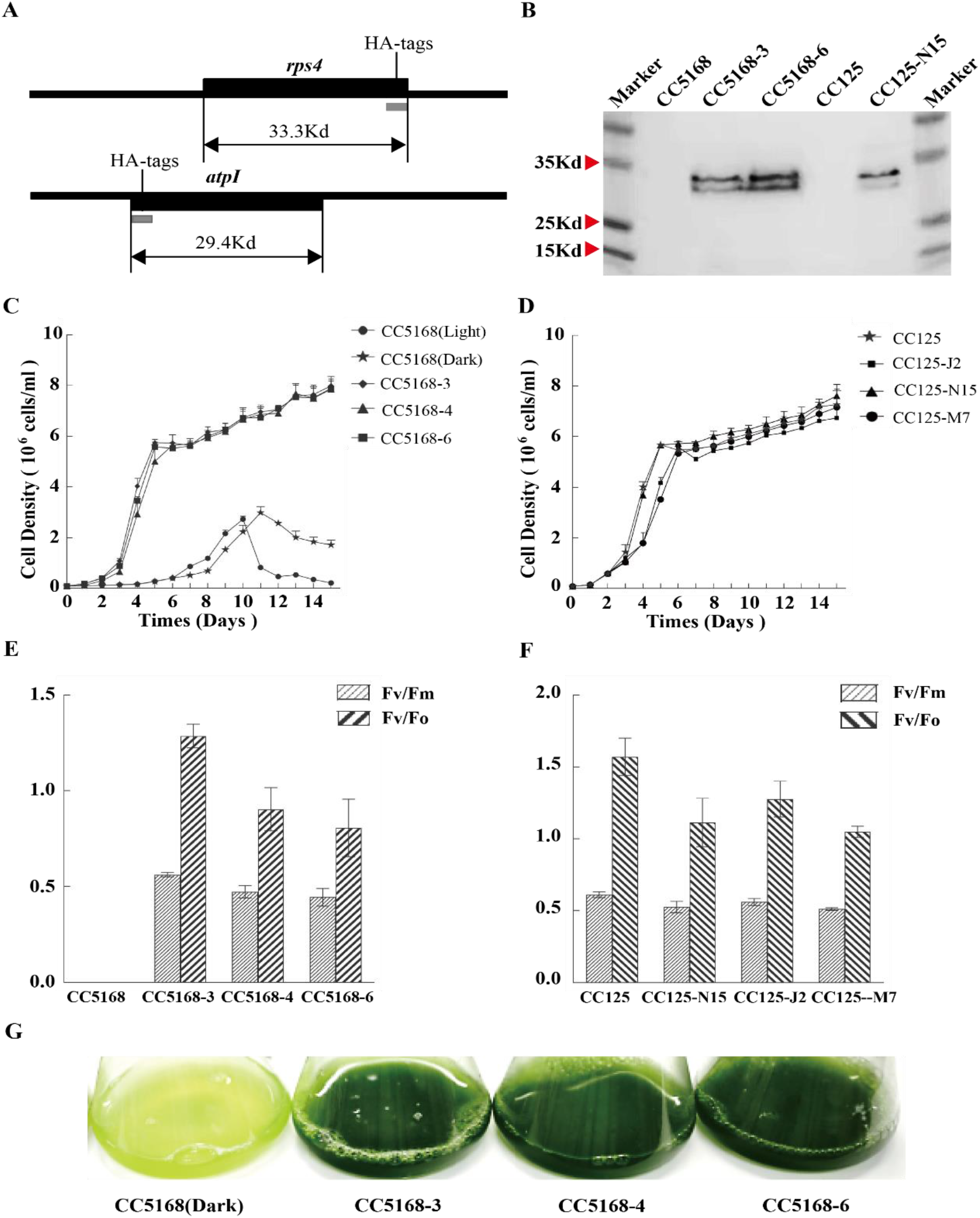
Functional analysis of transplastomic strains carrying SynCpV1.0. (A) HA-tags were incorporated into SynCpV1.0. The HA-tags were inserted in rps4 and atpI proteins, which were 33.3 Kd and 29.4 Kd, respectively. (B)Western blot analysis of transplastomic strains carrying SynCpV1.0. Total soluble protein was separated on Sodium Dodecyl Sulfate Polyacrylamide Gel Electrophoresis, transferred to polyvinyl chloride film membranes and probed with anti-HA antibody. CC5168 and CC125 were negative controls, and CC5168-3, CC5168-6 and CC125-N15 were transplastomic strains. (C)The growth curve of CC5168 and its transplastomic strains including CC5168-3, CC5168-4 and CC5168-6. CC5168 was cultured in light and dark conditions, respectively, as controls. (D) The growth curve of CC125 and its transplastomic strains including CC125-N5, CC125-J2, CC125-M7. Here, CC125 was applied as control. (E) The photosynthetic efficiency analysis of CC5168 and its transplastomic strains. CC5168-3, CC5168-4, CC5168-6 were the transplastomic strains harboring SynCpV1.0. (F) The photosynthetic efficiency analysis of CC125 and its transplastomic strains. CC125-N5, CC125-J2, CC125-M7 were the transplastomic strains harboring SynCpV1.0. (G) The SynCpV1.0 rescued the photosynthetic defect in mutant strain CC5168. CC5168-3, CC5168-4 and CC5168-6 were transplastomic strains originated from *psbH*-deficient mutant CC5168 which could not growth normally under light, and grow slowly without light.

We analyzed the photosynthetic efficiency of *C. reinhardtii* CC125 and *C. reinhardtii* CC5168, as well as their transplastomic strains to investigate the photosynthetic capacity of SynCpV1.0-harboring transplastomic strains. The results showed that *C. reinhardtii* CC5168 was unable to carry out photosynthesis activities at 22 °C under continuous light of 30 μmol m^−2^ s^−1^, whereas its transplastomic strains, such as CC5168-2, 5168-3, 5168-4, and 5168-6 performed normally photosynthesis activities, suggesting that the photosynthetic capacity of *C. reinhardtii* CC5168 was restored. We also found that the photosynthetic efficiency between *C. reinhardtii* CC125 and its transplastomic strains indicated no difference meaning that SynCpV1.0 exhibits the same biological functions as the wild-type cpDNA (Figure 5E, F and G).

To detect the transcriptional and translational activities of SynCpV1.0, we incorporated HA-tags into genes encoding rps4 (33.3 Kd) and atpI (29.4 Kd) proteins in SynCp V1.0. The total proteins of *C. reinhardtii* CC125 and its transplastomic strain CC125-N15, as well as *C. reinhardtii* CC5168 and its transplastomic strains (CC5168-3 and CC5168-6) were extracted and then subjected to western blot analysis with anti-HA-tag antibody. The experimental results showed that there was no protein band observed for *C. reinhardtii* CC5168 and CC125, while the protein bands of rps4 (33.3 Kd) and atpI (29.4 Kd) were detected from their transplastomic strains CC5168-3, CC5168-6, and CC125-N15 (Figure 5A and B), suggesting that the fully synthetic SynCpV1.0 genome can express the labeled functional proteins normally, i.e., the fully synthetic chloroplast genome exhibits the activities at the transcriptional and translational levels.

## DISCUSSION

The unicellular microalga *C. reinhardtii*, a model organism for photosynthesis research, has a single, easily transformable chloroplast and short cell-cycle. The molecular tools and techniques for the chloroplast engineering of *C. reinhardtii* have improved steadily since the first genetic transformation of chloroplast in 1988 (Boynton et al., 1988; Gimpel et al., 2016; Larrea-Alvarez and Purton, 2020; Macedo-Osorio et al., 2018). Hence, *C. reinhardtii* is the most desirable and novel platform for designing, modifying, replacing, and functional characterizing of the plastid genomes. In 2012, an artificial chloroplast genome with several modifications in function genes was assembled from the genome libraries of *C. reinhardtii* and transformed into the chloroplast (O’Neill et al., 2012). However, it remains unclear whether the artificial genome is step-wisely intergraded into the cpDNA by homologous recombination, or dose it directly replaced the wild cpDNA. Can we actually perform “replacing surgery” of chloroplast genome on photosynthetic organisms? We have not found further answers in the past 10 years. Here, we are presenting the first study on the *de novo* chemical synthesis, exogenous assembly, and functional characterization of an artificial cpDNA in photosynthetic organism.

The total synthesis and assembly of genomes serve as the basis and idle platform for subsequent in-depth functional studies and make a significant breakthrough in recent years, including the total synthesis and assembly of the genome of *Mycoplasma genitalium*(Gibson et al., 2008), and the Synthetic Yeast Genome Project-Sc2.0 (Mitchell et al., 2017; Richardson et al., 2017; Sanders et al., 2022). Undoubtedly, *de novo* chemical synthesis and functional assessment of cpDNAs in *C. reinhardtii* remains stagnant. The first hurdle is a large number of short dispersed repeats (SDRs) in the cpDNAs. About 20% of the chloroplast genome of *C. reinhardtii* is repetitive DNA containing various types of SDRs in the intergenic regions (Maul et al., 2002), which posed a great challenge to the synthesis and assembly of the chloroplast genome. For example, the chemical synthesis of the primary fragment seg 2, which contains more than 20% SDRs, failed. We eventually assembled the primary fragment seg 2 in yeast after optimizing and laborious screening (Supplementary figure S3). Therefore, the synthesis and assembly of DNA fragments similar to seg 2 should be avoided to reduce the workload in the subsequent redesign and modification of cpDNAs. Another hurdle is the homoplasmy of synthesis cpDNA in *C. reinhardtii* because of multiple copies of the cpDNA in *C. reinhardtii* (Chiang and Sueoka, 1967). In this study, SynCpV1.0 was transformed into the chloroplasts of *C. reinhardtii* and screened by gradient concentrations of streptomycin to reach a high degree of homoplasmy. The concentration of streptomycin used to achieve homogenization of SynCpV1.0 in this study was 4 times that of the transformation selection, and the transplastomic strains were subcultured with antibiotic selection pressure for more than 6 months.

Curiously, SynCpV1.0 owned the characteristics of a shuttle vector showing that SynCpV1.0 from transplastomic algae could re-transformed into *E. coli*, which suggested that circular SynCpV1.0 is present in the chloroplast genome of *C. reinhardtii*. Considering that SyncpV1.0 was assembly in yeast, function characterized in the chloroplast of *C. reinhardtii*, and re-transformed into *E. coli*, we believe that the SynCpV1.0 can shuttle among the *C. reinhardtii’s* chloroplasts, yeast and *E. coli*. This would be an astonishing discovery for plastid genomes, which are difficult to edit and modify. Currently, the commonly used gene-editing tools, e.g.,CRISPR-Cas 9 or TALEN have slow progress and low efficiency in editing the plastid genomes. Our study has determined for the first time that the fully synthetic cpDNA has the characteristics of a shuttle vector. We can take advantage of the well-established gene-editing technique for *E. coli* to edit and rectify the fully synthetic cpDNA, providing a new avenue for redesigning and modifying cpDNA based on SynCpV1.0 in the future. Through transformation and antibiotic selection, we obtained transplastomic algal strains that are homoplasmic for SynCp V1.0, which was generated *via de novo* chemical synthesis and *in vitro* assembly. In addition, our results from this study confirmed that the synthetic cpDNA owns a great potential to perform biological activities. Hence, our study has opened a new chapter for the deep investigation of chloroplast of algae and other photosynthetic plants, allowing the comprehensive studies on cpDNAs and exploration of synthetic biology at the subcellular level *via* the bottom-up recoding and total synthesis of cpDNAs. Moreover, our study has also laid the foundation for subsequent functional studies of artificial organelles and redesigning the photosynthetic system in microalgae.

## MATERIALS AND METHODS

### Cultivation of algal, bacterial, and yeast strains

The strains of *C. reinhardtii*, which are used for transformation in chloroplast were obtained from the *Chlamydomonas* Resource Center (St. Paul, MN 55108, USA). A cell wall-deficient strain of *C.reinhardtii* CC5168, which is unable to perform the function of *psbH* gene and non-photosynthetic until transformed, was maintained in TAP medium under dim light (5–10 μE m^−2^ s^−1^) in order to avoid from any light damage (Wannathong et al., 2016). The *C. reinhardtii* CC125 (mt+) was a wild-type strain, which was grown under standard illumination (40–50 μE m^−2^ s^−1^) at 25 °C in a TAP medium, as previously described (Harris et al., 1989; Kropat et al., 2011). Streptomycin antibiotic (150 mg/L) or Paromomycin (10 mg/L) was added to TAP agar medium as required. To reach homoplasmy for SynCpV1.0 in *C. reinhardtii* chloroplasts, the transplastomic algal strains were exposed to progressively increasing selection pressure of streptomycin within the concentration range of 150–600 mg/L.

*S. cerevisiae* strains, including JDY19 (Mat a, thr4 Mal’), JDY20 (Mat alpha, thr4 Mal’), BY4741 (Mata, his3Δ1 leu2Δ0 ura3Δ0 met15Δ0), and HWY175 (Mat alpha, met17Δ::KanMX4 his3Δ1 leu2Δ0 lys2Δ0 ura3Δ0) were provided by Professor Dai (Luo et al., 2021; Schindler et al., 2018). The strains of *S. cerevisiae* were cultivated in Yeast Extract Peptone Dextrose Adenine medium (1% yeast extract, 2% peptone, 0.01% adenine hemisulfate, and 2% glucose) or in synthetic complete (SC) dropout medium (0.17% Difco yeast nitrogen base without amino acids and ammonium sulfate, 0.5% ammonium sulfate and 0.083% amino-acid dropout mix, 0.01% adenine hemisulfate and 2% glucose) (Sherman, 2002). The strains of *S. cerevisiae* were cultured at 30 °C and at 250 rpm rotation in baffled shake-flasks. *E. coli* EPI300 strains were cultured at 37 °C and at 250 rpm in the Luria broth medium (Fisher Scientific, Pittsburgh, PA, USA).

### *De novo* synthesis of SynCpV1.0

SynCpV1.0, which has a total length of 221.372 bp, was divided into 44 primary fragments (seg1–44), each with a length of 4.915 kb except for seg44 (4.901 kb). All primary fragments were synthesized by Qinglan Biotech Co., Ltd (Wuxi, China). All primary fragments were cloned into the pUC18 vector, from which each primary fragment can be obtained *via* cleavage caused by the restriction endonuclease *Not* I. The sequences of these primary fragments were confirmed *via* DNA sequencing and restriction endonuclease digestion.

The assembly of the primary fragment seg2: The linear pRS416 plasmid fragment, seg-a, seg-b1, seg-b2, seg-c, and seg-d were co-transformed into *S. cerevisiae* BY4741, followed by the screening for yeast transformants with the SC-URA selective medium to obtain seg2. The assembly of the secondary fragment chunk1: the linear BAC vector fragment, DNA fragment of the selectable marker gene, 7-35 junction fragment, seg35-seg44, seg1-seg7 were co-transformed into *S. cerevisiae* BY4741, followed by the screening for yeast transformants with the SC-URA selective medium. The resulting secondary fragment chunk1 was validated *via* junction PCR with 17 primer pairs (Supplementary table S1). The assembly of the secondary fragment chunk2: the linear pRS415 vector fragment and the primary fragments seg7-21 were co-transformed into *S. cerevisiae* HWY175, and the resulting yeast transformants were screened with the SC-LEU selective medium. The resulting secondary fragment chunk2 was validated *via* junction PCR with 17 primer pairs (Supplementary table S1). The assembly of the secondary fragment chunk3: the linear pRS411 vector fragment and the primary fragments seg22-35 were co-transformed into *S. cerevisiae* BY4741, which was then screened for yeast transformants with the SC-MET selective medium. The resulting secondary fragment chunk3 was validated *via* junction PCR with 14 primer pairs (Supplementary Table S1). The yeast transformation and the extraction of yeast genomic DNA were carried out according to methods described by Gietz (Gietz and Schiestl, 2007) and Gibson (Gibson, 2009) respectively.

The yeast strains carrying chunk2 and chunk3 were crossed and screened using the SC-LEU-MET agar plate for diploid hybrids harboring chunk2 and chunk3. The selected diploid yeast strains were subject to the induction of sporulation and the separation of spores to obtain the haploid yeast cells, which were subsequently screened with SC-LEU and SC-MET agar plates for haploid yeast strains containing chunk2 and chunk3. After being crossed with *S. cerevisiae* JDY19 and JDY20, the resulting haploid yeast strains of the mating-type alpha were selected for hybridization with the chunk1-harboring yeast strain, followed by the screening with the SC-LEU-MET-URA agar plate for the yeast strain containing all three secondary fragments. The yeast strain obtained in the previous step was transformed with the ZLP012 plasmid (harboring genes encoding for the endonuclease *I-SceI* and the auxotrophic selectable marker HIS3) and screened for the yeast transformant carrying chunk1, chunk2, chunk3, and ZLP0121 using the SC-URA-LEU-MET-HIS agar plate. Subsequently, the yeast transformant was treated with galactose to induce the expression of *I-SceI*, which cleaved and linearized chunk1, chunk2, and chunk3 at the *I-SceI* restriction sites. Those three linearized DNA fragments were assembled in yeast cells into a complete cpDNA, which was screened and validated *via* junction PCR with primers (Supplementary material Table S2) and DNA sequencing, respectively.

### Genetic transformation of *C. reinhardtii* chloroplasts

The SynCpV1.0-harboring plasmid was extracted from yeast in accordance with a previously described method (Gibson, 2009) and transformed into *E. coli* EPI300 *via* electroporation (Chen et al., 2006). Then, SynCpV1.0 was isolated from the *E. coli* transformant using the QIAGEN Plasmid Midi Kit (REF.12143, Hilden, Germany) according to instructions provided by the manufacturer.

After that, the fully synthetic SynCpV1.0 plasmid was transformed into chloroplast of *C. reinhardtii* with the biolistic method described by Zapata et al. with slight modifications (Guzmán-Zapata et al., 2016). Briefly, the recipient algal cells with a cell density of 2 ×10^6^ cell/mL were harvested by centrifugation at 3500 ×g for 5 min. The cell pellet was resuspended with fresh TAP medium to a cell density of 1 × 10^8^ cell/mL. Then, 250 μL of the algal cell suspension was spread at the center of a selective agar plate (90 × 15 mm) and incubated under a dry condition in the dark at 25 °C for 2 h.

Preparation of DNA-coated microparticles for bombardment: A total of 50 μL of microparticles was sequentially added with 5.0 μL of DNA (1.0 μg/μL), 50 μL of 2.5 M CaCl_2_, and 20 μL of 0.1 mM spermidine, followed by a thorough mixing *via* vortex. After centrifugation at 10000 rpm for 1 min, the resulting pellet was rinsed with 70% alcohol and resuspended with 100 μL of absolute ethanol. A total of 10 μL of the DNA-coated microparticle solution was used for each bombardment. The bombardment was carried out by the Biolistic PDS-1000/He Particle Delivery System (BIO-RAD, California, USA) at a rupture disc under pressure of 1,100 psi. The bombardment was carried out at a distance of 9 cm. After the bombardment, the plate was incubated in the dark at 25 °C for 24 h, followed by incubation at 25 °C in an incubator with a photoperiod ratio of 16:8 (day: night) for 3–4 weeks until single green colonies were observed.

### Screening of transplastomic algal strains harboring SynCpV1.0

Templates for PCR analysis were prepared by suspending cells in 50 μL of 5% Chelex-100 (w/v), after that they were incubating at 98 °C for 30 min following by cooling to 10 °C and then 2 μL of lysate was used for each PCR reaction. Transplastomic strains were subjected to the extraction of genomic DNA, which were then used as templates for PCR assay using the primer pairs aphVIII-F/R, BAC44-F/R, BAC1-F/R etc. (Supplementary Table S3) to further screen for transplastomic strains harboring all the exogenous fragments.

To achieve the homoplasmy for SynCpV1.0 in *C. reinhardtii* chloroplast, the transplastomic *C. reinhardtii* strains were exposed to progressively increasing selection pressure of streptomycin within the concentration range of 150–600 mg/L, and subcultured with antibiotic selection pressure for more than 6 months.

### RFLP-southern blotting analysis

The total DNA of the transplastomic algal strains harboring SynCpV1.0 was isolated by the M5 Hiper Plant Genomic DNA Max Kit (Cat. MF136-01, Mei5 Biotechnology Co. Ltd, Beijing, China) for RFLP-Southern blotting analysis. Firstly, 10 μg of total genomic DNA of algae were digested with *Hin*d III or *Eco*R V endonuclease enzymes for 6 h at 37 °C in a total volume of 50 μL, then separated by 0.8% agarose gel electrophoresis, finally transferred to a Nylon membrane (Cat. FFN13, Beyotime, Shanghai, China) (Khandjian and E., 1987; Southern, 1992). The DIG High Prime DNA Labeling and Detection Starter Kit II (Roche, Indianapolis, IN, USA) was used to conduct southern blots according to the manufacturer’s protocol. Probes specific for sequences adjacent to integration sites for *aphVIII* and *psaA* were generated using primers aphVIII-F1/R1 and psaA-F/R PCR (Supplementary Table S4), respectively.

### Re-transformation of SynCpV1.0 from transplastomic algal strains into *E. coli*

The total genomic DNA from transplastomic strains harboring SynCpV1.0 and the recipient algal strain CC125 was extracted with M5 Hiper Plant Genomic DNA Max Kit (Cat. MF136-01, Mei5 Biotechnology Co. Ltd, Beijing, China). Then about 10 μg of total genomic DNA was transformed into 100 μL competent cells of *E. coli* EPI300 according to the conventional electroporation method (*51*). Plasmids extracted from those positive colonies with the QIAGEN Plasmid Midi Kit (REF.12143, Hilden, Germany), were used as templates for the PCR amplification of target sequences on *aphVIII*, BAC vector backbone and the fragments among the adjacent primary fragments *via* junction PCR (Supplementary Table S2). The resulting PCR products were separated with 1.2% agarose gel electrophoresis. Those positive colonies were conducted DNA sequencing by Wuhan Frasergen Bioinformatics Co., Ltd.

### Western blot analysis

The total protein (5–015μg) from the transplastomic algal strains harboring SynCpV1.0 was extracted and subjected to 12% or 15% (v/v) Sodium Dodecyl Sulfate Polyacrylamide Gel Electrophoresis in order to perform immunoblot analyses and then transferred to polyvinyl chloride film membranes (Hirano, 2012). Then the membranes were successively blocked in TBS-T (20 mM Tris, 137 mM NaCl, 1 M HCl pH 7.4, 0.1% Tween-20) with 0.5% milk powder overnight at 4 °C, incubated with a primary antibody in TBS-T with 0.5% milk at room temperature for 1 h, washed in TBS-T 3 times and for 5 min each time, incubated with secondary antibody in TBS-T with 0.5% milk (1 h at room temperature), and finally washed again in TBS-T as above. The probed blots were scanned and quantified with an Odyssey Infrared Imaging System (LI-COR, Lincoln, NE). The primary antibodies and dilutions were used as follows: anti-hemagglutinin (HA) antibody (ab18181, Abcam, Cambridge, MA, USA; 1: 1000). Secondary anti-rat (ab205719, Abcam, Cambridge, MA, USA) antibodies were diluted to 1:5000.

### Determination of photosynthetic efficiency in *C. reinhardtii*

The algal cells in the logarithmic phase were inoculated into fresh TAP medium at an initial cell density of 4 × 10^4^ cells/mL, and the growth was monitored by determining the cell density every day with a cell counter until entering the stationary phase. The growth curves of different algal strains were plotted with cell density as the y-axis and time as the x-axis. The Fv/Fm, Fv/Fo, and ΔF/Fm of algal strains were measured by the Phyto-PAM Chlorophyll Fluorometer (Zealquest Scientific Technology Co., Ltd., Shanghai, China). Measurements were performed three times and the averages were calculated.

### Statistical analyses

All statistical analyses, including arithmetic means and standard deviations, and parametric two-tailed Student’s tests, were applied in all cases.

## ACKNOWLEDGMENTS

We thank Prof. Junbiao Dai for providing yeast strains including JDY19, JDY20, BY4741 and HWY175. We also thank Prof. Zhouqing Luo for his crucial technical guidance in the synthesis and assembly of SynCpV1.0. This work was supported by Chinese National Key R & D Project for Synthetic Biology (2018YFA0902500), National Natural Science Foundation of China (32273118, 31870343, 31970366), Shenzhen Basic Research Projects (JCYJ20180507182405562) and Shenzhen Special Fund for Sustainable Development (KCXFZ20211020164013021).

## AUTHOR CONTRIBUTIONS

Z.L.H., and C.G.W. conceived the project. C.L.G., G.Y.Z., H.W., R.M., and X.Y.L. generated and analyzed data and generated figures. Z.L.H., C.G.W., and B. J. provided funding and project management. C.L.G., G.Y.Z., B.J.,C.G.W., and Z.L.H. wrote the paper. Z.L.H., C.G.W., B.J., and H. L. reviewed and edited the paper. All other authors contributed edits and comments.

## DECLARATION OF INTERESTS

The authors declare no competing interests.

## Notes

### Competing Interest Statement

The authors have declared no competing interest.

