## Supplementary Materials for "The “replacing surgery” of cpDNA: *de novo* chemical synthesis and *in vivo* functional testing of *Chlamydomonas* chloroplast genome"

**This PDF file includes:**

Tables S1, S2, S3 and S4

Figures. S1, S2 and S3

### Table S1: List of primers for secondary fragments chunk1, chunk2, and chunk3

| **Primers for secondary fragments chunk1** | |
|---|---|
| **Name** | **Sequence (5'-3')** |
| BAC2 | ACTATTAGGCTATGACTGGGC |
| V-R | CGGGTACGAGGAAAGTGCAA |
| 1-F | CGAAGGGGACGTATCCGAAAT |
| 1-R | ACCCATAGCAGCTGGGTCTA |
| 2-F | TTGCAAACACGATGCGGATT |
| 2-R | CCCGCTAGCGCAGTATTTCA |
| 3-F | GAGGATTTGCAATCCACGGC |
| 3-R | TTAGAGGCGCTGCTTCAACT |
| 4-F | AACCGCCCAGTAACCAACTT |
| 4-R | ACCGTCCAACAGTAGCAGAAG |
| 5-F | GCAAAGGCCCGTTTAAGGTG |
| 5-R | GCAATGCGAGCATTACGGTT |
| 6-F | CGTCTCCTTACGGGGACATTT |
| 6-R | AGTTACACCGCCTACTGCTTT |
| 7-F | CAGCAGCACGCCAACTTATT |
| 34-R | CCTGCGGTTGACCCATTAGA |
| 35-F | TGCTTTGTACGCCACCATTG |
| 35-R | TGTACGGAGCAGATGGAGAGA |
| 36-F | ATTCCCTCGGAATGGCAGTG |
| 36-R | TTCCAGGTGTTGGCCACAAT |
| 37-F | ATATGCCTCGGGCATCCAAG |
| 37-R | GTCGTTGATCAAAGCTCGCC |
| 38-F | GGGACGTCCTAGGCAAATGA |
| 38-R | TGTCCGCTTCATCAGACACG |
| 39-F | AGGGTATTTCTCGGTTGGGA |
| 39-R | CTGTGCTAGCTGGTCCTAGTG |
| 40-F | GGCTGCACGATAGCGTAAAC |
| 40-R | GTCACGGCCACCTACTGTAA |
| 41-F | TGACCACACAGCGTAAAGCA |
| 41-R | CAACCCCGAAGGGGAGTAAC |
| 42-F | ACCAGCATTTGCTTCGGCTA |
| 42-R | ATTGTGGTGGGACCACGTTT |
| 43-F | GAAAGAGCGAGCTGTGCTTG |
| 43-R | GTAGCCCCGCAGTCGTTTAT |
| 44-F | TAAAAATGCCATCCGCCCCA |
| BAC1 | AGCCAGTGTGATTACTCGTCC |

| **Primers for secondary fragments chunk2** | |
|---|---|
| **Name** | **Sequence (5'-3')** |
| M13-F | ACACAGGAAACAGCTATGAC |
| 6-R | AGTTACACCGCCTACTGCTTT |
| 7-F | CAGCAGCACGCCAACTTATT |
| 34-F | TGCTTTGTACGCCACCATTG |
| 8-F | ACTGCCGGTAGAGACACTAA |
| 8-R | GCTCTCACGAGATCGCTTCT |
| 9-F | TGGTGAGAATCCAATGCCCC |
| 9-R | CATTACTGCAAGCACACGCA |
| 10-F | CCAACAGGCTGTCACGAAGT |
| 10-R | GCGTAACGCTCACAACTTCC |
| 11-F | AGAACGAGTGCTGACGTGTT |
| 11-R | GGGCAACTATCGTTCCACCA |
| 12-F | AAGGAGGCAGCGTTCCTTTT |
| 12-R | CCCTCTTTGGGACGTCCTTC |
| 13-F | TCGGAAGGAGAATGTTGCCC |
| 13-R | ATCAAGGCAGCAACCTGTGT |
| 14-F | GCCTGGCTTTAGGGTCAGTT |
| 14-R | TGGGCTGTTGGGTGTTTCTT |
| 15-F | TGTAAACCTGCTCGTGCCAT |
| 15-R | GAGAAGCACTGCACACCGTA |
| 16-F | GCAACTTTTTGTTGCGGTGC |
| 16-R | AGTGCCACGGCTTACTTCAA |
| 17-F | CTGGTCGGGCAAGTAAACCT |
| 17-R | GCTCCACAGCAGCAAAGTTG |
| 18-F | TGGTGGAATGCTAACAACCGT |
| 18-R | GAAGGGGACGCTCTTTCACT |
| 19-F | TTCACCATAATGCATGCCGC |
| 19-R | ACACCAGGTGCTACGTTAGA |
| 20-F | GGCCCATTAAACCACCACCT |
| 20-R | TGAGCAACTGGCACTAGTCT |
| 21-F | TTCAAGGCTGCGTCACATCT |
| 21-R | AGGCTTAGAAGCACGAGCAG |
| 22-F | GAACTGCCTGCAGCTTTTGG |
| M13-R | CGCCAGGGTTTTCCCAGTCACGAC |
| **Primers for secondary fragments chunk3** | |
| **Name** | **Sequence (5'-3')** |
| M13-F | CGCCAGGGTTTTCCCAGTCACGAC |
| 21-R | AGGCTTAGAAGCACGAGCAG |
| 22-F | GAACTGCCTGCAGCTTTTGG |
| 22-R | AGAGCTAACAAATCCCCGGC |
| 23-F | TGAACGGGCAGTTTGCCATA |

| | |
|---|---|
| 23-R | GCACTCCCGGAGGAACTAGA |
| 24-F | TCGTAACCGCACCCGTAAAA |
| 24-R | GGCAGTTGGCAGGGGATTTA |
| 25-F | GATCGAGCATCTCGTGAGACA |
| 25-R | GAGCTGCACGACAAACCTTG |
| 26-F | ATGGGACTCGAACCCACAAC |
| 26-R | TGGTGTTCAATGCCAGGTGT |
| 27-F | AATGACCCTGCTCGTAACCC |
| 27-R | TGCGCTGTACATCGACTTGG |
| 28-F | GCCATGGTGGCTTTGGTTTT |
| 28-R | GCAGACCTTAAAGGCAGGGA |
| 29-F | TGCCCGATTAGCTGTGTAGC |
| 29-R | GCAGATGGAAGCCATCCAGA |
| 11-F | AGAACGAGTGCTGACGTGTT |
| 11-R | GGGCAACTATCGTTCCACCA |
| 10-F | CCAACAGGCTGTCACGAAGT |
| 10-R | GCGTAACGCTCACAACTTCC |
| 9-F | TGGTGAGAATCCAATGCCCC |
| 9-R | CATTACTGCAAGCACACGCA |
| 8-F | ACTGCCGGTAGAGACACTAA |
| 8-R | GCTCTCACGAGATCGCTTCT |
| 34-F | GTCCTGAAGGGAAAGGTGCAA |
| 34-R | CCTGCGGTTGACCCATTAGA |
| 35-F | TGCTTTGTACGCCACCATTG |
| M13-R | ACACAGGAAACAGCTATGAC |

**Table S2: List of primers for identifying the yeast/*E.coli* strain containing the complete SynCp V1.0 genome**

| **Name** | **Sequence (5'-3')** |
|---|---|
| BAC2 | ACTATTAGGCTATGACTGGGC |
| V-R | CGGGTACGAGGAAAGTGCAA |
| 1-F | CGAAGGGGACGTATCCGAAAT |
| 1-R | ACCCATAGCAGCTGGGTCTA |
| 2-F | TTGCAAACACGATGCGGATT |
| 2-R | CCCGCTAGCGCAGTATTTCA |
| 3-F | GAGGATTTGCAATCCACGGC |
| 3-R | TTAGAGGCGCTGCTTCAACT |
| 4-F | AACCGCCCAGTAACCAACTT |
| 4-R | ACCGTCCAACAGTAGCAGAAG |

| | |
|---|---|
| 5-F | GCAAAGGCCCGTTTAAGGTG |
| 5-R | GCAATGCGAGCATTACGGTT |
| 6-F | CGTCTCCTTACGGGGACATTT |
| 6-R | AGTTACACCGCCTACTGCTTT |
| 7-F | CAGCAGCACGCCAACTTATT |
| 34-F | GTCCTGAAGGGAAAGGTGCAA |
| 8-F | ACTGCCGGTAGAGACACTAA |
| 8-R | GCTCTCACGAGATCGCTTCT |
| 9-F | TGGTGAGAATCCAATGCCCC |
| 9-R | CATTACTGCAAGCACACGCA |
| 10-F | CCAACAGGCTGTCACGAAGT |
| 10-R | GCGTAACGCTCACAACTTCC |
| 11-F | AGAACGAGTGCTGACGTGTT |
| 11-R | GGGCAACTATCGTTCCACCA |
| 12-F | AAGGAGGCAGCGTTCCTTTT |
| 12-R | CCCTCTTTGGGACGTCCTTC |
| 13-F | TCGGAAGGAGAATGTTGCCC |
| 13-R | ATCAAGGCAGCAACCTGTGT |
| 14-F | GCCTGGCTTTAGGGTCAGTT |
| 14-R | TGGGCTGTTGGGTGTTTCTT |
| 15-F | TGTAAACCTGCTCGTGCCAT |
| 15-R | GAGAAGCACTGCACACCGTA |
| 16-F | GCAACTTTTTGTTGCGGTGC |
| 16-R | AGTGCCACGGCTTACTTCAA |
| 17-F | CTGGTCGGGCAAGTAAACCT |
| 17-R | GCTCCACAGCAGCAAAGTTG |
| 18-F | TGGTGGAATGCTAACAACCGT |
| 18-R | GAAGGGGACGCTCTTTCACT |
| 19-F | TTCACCATAATGCATGCCGC |
| 19-R | ACACCAGGTGCTACGTTAGA |
| 20-F | GGCCCATTAAACCACCACCT |
| 20-R | TGAGCAACTGGCACTAGTCT |
| 21-F | TTCAAGGCTGCGTCACATCT |
| 21-R | AGGCTTAGAAGCACGAGCAG |
| 22-F | GAACTGCCTGCAGCTTTTGG |
| 22-R | AGAGCTAACAAATCCCCGGC |
| 23-F | TGAACGGGCAGTTTGCCATA |
| 23-R | GCACTCCCGGAGGAACTAGA |
| 24-F | TCGTAACCGCACCCGTAAAA |
| 24-R | GGCAGTTGGCAGGGGATTTA |
| 25-F | GATCGAGCATCTCGTGAGACA |
| 25-R | GAGCTGCACGACAAACCTTG |
| 26-F | ATGGGACTCGAACCCACAAC |

| | |
|---|---|
| 26-R | TGGTGTTCAATGCCAGGTGT |
| 27-F | AATGACCCTGCTCGTAACCC |
| 27-R | TGCGCTGTACATCGACTTGG |
| 28-F | GCCATGGTGGCTTTGGTTTT |
| 28-R | GCAGACCTTAAAGGCAGGGA |
| 29-F | TGCCCGATTAGCTGTGTAGC |
| 29-R | GCAGATGGAAGCCATCCAGA |
| 11-F | AGAACGAGTGCTGACGTGTT |
| 11-R | GGGCAACTATCGTTCCACCA |
| 10-F | CCAACAGGCTGTCACGAAGT |
| 10-R | GCGTAACGCTCACAACTTCC |
| 9-F | TGGTGAGAATCCAATGCCCC |
| 9-R | CATTACTGCAAGCACACGCA |
| 8-F | ACTGCCGGTAGAGACACTAA |
| 8-R | GCTCTCACGAGATCGCTTCT |
| 34-F | GTCCTGAAGGGAAAGGTGCAA |
| 34-R | CCTGCGGTTGACCCATTAGA |
| 35-F | TGCTTTGTACGCCACCATTG |
| 35-R | TGTACGGAGCAGATGGAGAGA |
| 36-F | ATTCCCTCGGAATGGCAGTG |
| 36-R | TTCCAGGTGTTGGCCACAAT |
| 37-F | ATATGCCTCGGGCATCCAAG |
| 37-R | GTCGTTGATCAAAGCTCGCC |
| 38-F | GGGACGTCCTAGGCAAATGA |
| 38-R | TGTCCGCTTCATCAGACACG |
| 39-F | AGGGTATTTCTCGGTTGGGA |
| 39-R | CTGTGCTAGCTGGTCCTAGTG |
| 40-F | GGCTGCACGATAGCGTAAAC |
| 40-R | GTCACGGCCACCTACTGTAA |
| 41-F | TGACCACACAGCGTAAAGCA |
| 41-R | CAACCCCGAAGGGGAGTAAC |
| 42-F | ACCAGCATTTGCTTCGGCTA |
| 42-R | ATTGTGGTGGGACCACGTTT |
| 43-F | GAAAGAGCGAGCTGTGCTTG |
| 43-R | GTAGCCCCGCAGTCGTTTAT |
| 44-F | TAAAAATGCCATCCGCCCCA |
| BAC1 | AGCCAGTGTGATTACTCGTCC |
| 34-35F1 | ATCATCTAATTCACCAGCGAAAAT |
| 34-35R1 | CTGCTCCTGCTACAACATTCG |
| 35-36F1 | TTCTACCCTTTCCCTAACGG |
| 35-36R1 | GCCGATAGGCGAGTTAGTAG |
| 41-42F1 | ATTACCCTTTCAGGCCGATAAA |
| 41-42R1 | CGATAGGCGAGGTAACTGCT |

*Primers 34-35F1/R1, 35-36F1/R1 and 41-42F1/R1 were used to validate the plasmid recovered in the *E.coli* EPI300, instead of 35F/R, 36F/R and 42F/R. In the list, 7-F together with 34 F were used to validate the adjacent fragments seg7-seg8. 11F/R, 10F/R, 9F/R and 8F/R after 29F/R were used to validate the adjacent fragments seg30-seg31, seg31-seg32, seg32-seg33 and seg33-seg34, respectively.

**Table S3: List of primers for screening transplastomic algal strains harboring SynCp V1.0**

| Name | Sequence (5'-3') |
|---|---|
| aphVIII-F | CCACCTACGGCAAGCTGAC |
| aphVIII-R | GAAGCCGATAAACACCAGCC |
| BAC44-F | TTTAGTGGCAGTTGCCTCCTT |
| BAC44-R | GGTAGTCGCCCTGCTTTCTC |
| BAC1-F | GAGGGTGGTTCGTCACATTT |
| BAC1-R | TACCGCCACTGCCTATGTT |

**Table S4: List of primers for probes specific for sequences adjacent to integration sites for *aphVIII* and *psaA***

| Name | Sequence (5'-3') |
|---|---|
| aphVIII-F1 | CCACCTACGGCAAGCTGAC |
| aphVIII-R1 | GAAGCCGATAAACACCAGCC |
| psaA-F | CAGAAGCTGTAACCGTACCCC |
| psaA-R | AGACCGTGTAATTCGTCACCG |

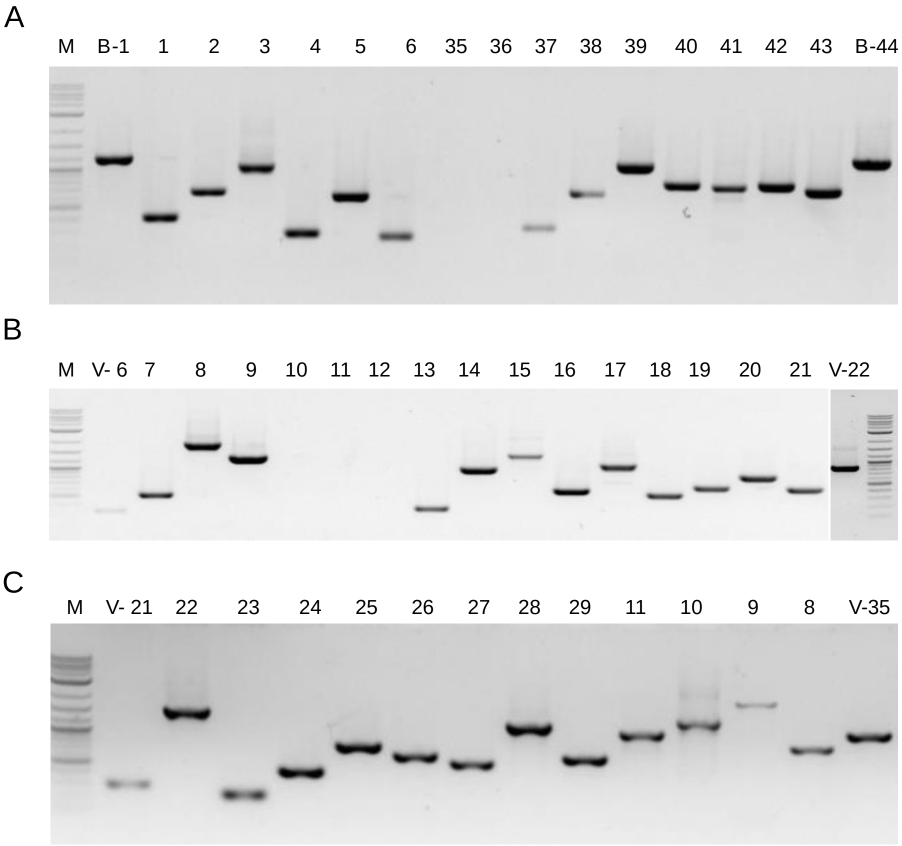


**Figure S1. Identify of the secondary fragments assembled in the *S. cerevisiae***

(A) Junction PCR of assembled chunk1. The primary fragments were assembled with the fragment of BAC in *S. cerevisiae* BY4741, B-1 represents the primers BAC2 and V-R generating junction sequences between vector BAC and seg1, B-44 represents the primers 44-F and BAC1, generating junction sequences between vector BAC and seg44. 1-6 and 35-43 are the junction primers that amplify the fragments of the two adjacent fragments.

(B) Junction PCR of assembled chunk2. The primary fragments and fragment of pRS415 vector were co-transformed into *S. cerevisiae* HWY175, V-6 represents the primers M13-F and 6-R generating junction sequences between vector pRS415 and seg6, and V-22 represents the primers 22-F and M13-R generating junction sequences between vector pRS415 and seg22, respectively. 7-22 are the junction primers that amplify the fragments of the two adjacent fragments.

(C) Junction PCR of assembled chunk3. The primary fragments and fragment of pRS411 vector were co-transformed into *S. cerevisiae* BY4741, V-21 represents the primers M13-F and 21-R generating junction sequences between vector pRS411 and seg21, and V-35 represents the primers 35-F and M13-R generating junction sequences between vector pRS411 and seg35, respectively. 22-29 and 11-8 are the junction primers that amplify the fragments of the two adjacent fragments. M in (A), (B) and (C) represents DL1,2000 marker.

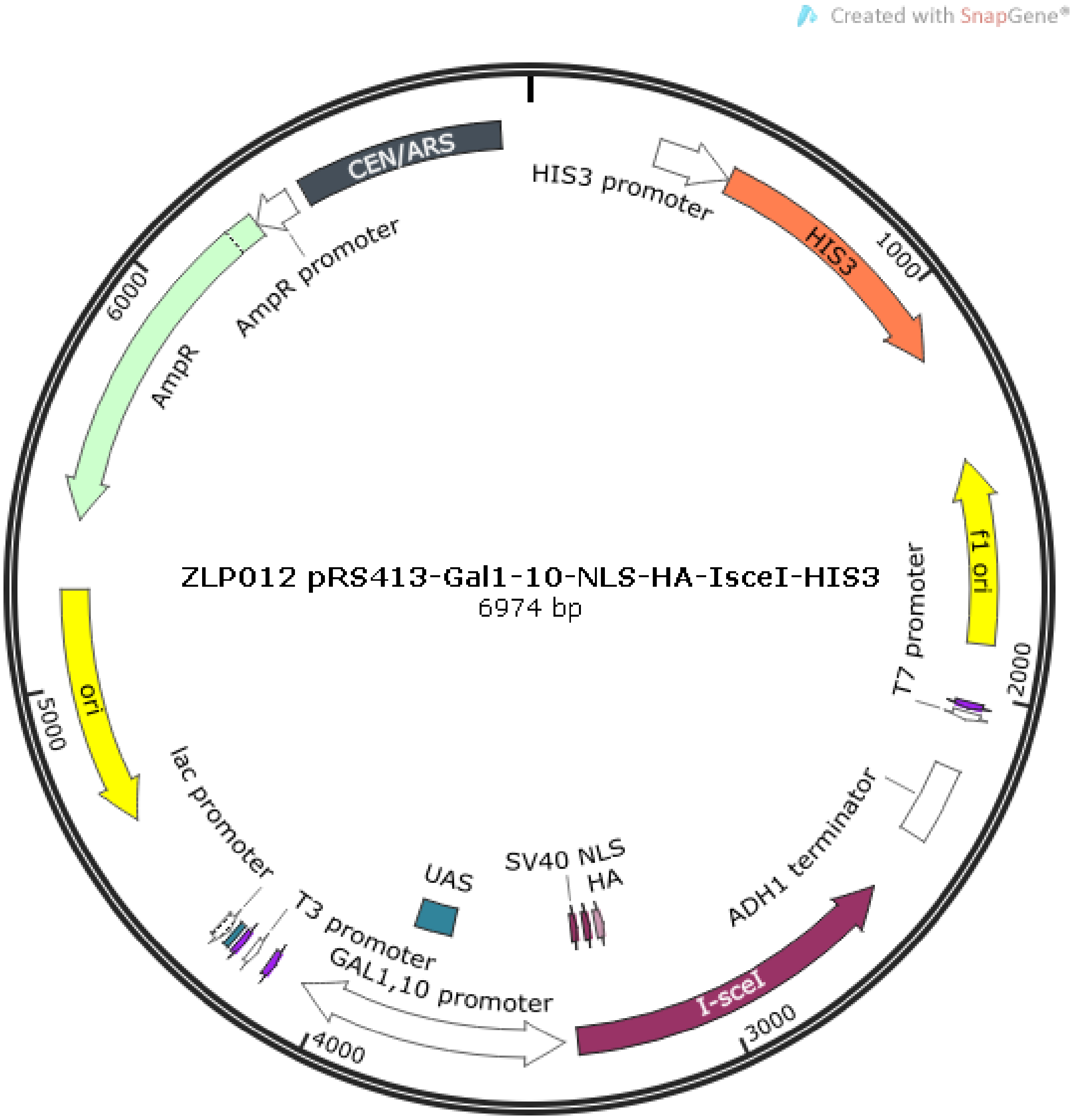


**Figure S2. The map of plasmid ZLP102**

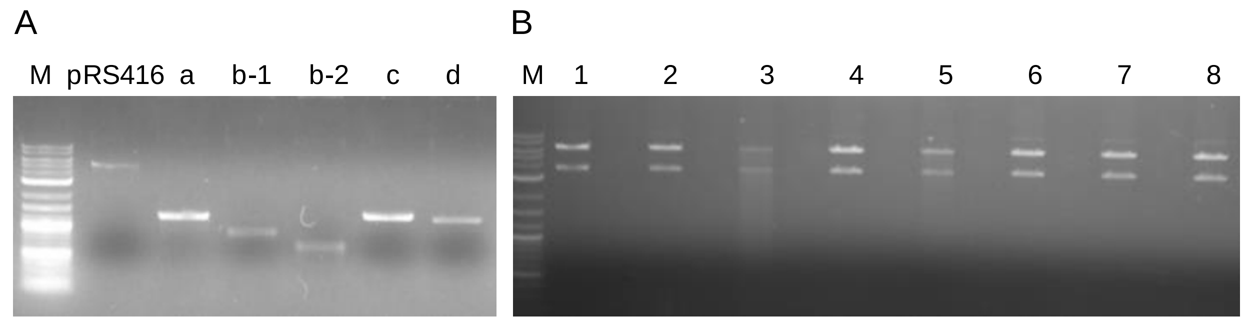


**Figure S3. The assemble of seg2**

(A) The primary fragments a, b-1, b-2, c, and d, and also, the vector fragment of pRS416. All the fragment were co-transformed into *S. cerevisiae* BY4741. M represents DL1,2000 marker.

(B) Restriction enzyme digestion verification. Lane 1-8 are the productions of enzyme digestion of plasmids extracted from the 8 positive transformants by SpeXia. M represents DL1,2000 marker.
